# Pre-diabetes increases tuberculosis disease severity, while high body fat without impaired glucose tolerance is protective

**DOI:** 10.1101/2021.04.06.438735

**Authors:** Roma Sinha, Minh Dao Ngo, Stacey Bartlett, Helle Bielefeldt-Ohmann, Sahar Keshvari, Sumaira Z. Hasnain, Meg L. Donovan, Jessica C. Kling, Antje Blumenthal, Chen Chen, Kirsty R. Short, Katharina Ronacher

## Abstract

Type 2 diabetes (T2D) is a well-known risk factor for tuberculosis (TB), but little is known about pre-diabetes and the relative contribution of impaired glucose tolerance *vs*. obesity towards susceptibility to TB. Here, we developed a preclinical model of pre-diabetes and TB. Mice fed a high fat diet (HFD) for 12 weeks presented with impaired glucose tolerance and hyperinsulinemia compared to mice fed normal chow diet (NCD). Infection with *M. tuberculosis* (Mtb) H_37_R_v_ after the onset of dysglycemia was associated with significantly increased lung pathology, lower concentrations of TNF-a, IFN-g, IFN-β and IL-10 and a trend towards higher bacterial burden at 3 weeks post infection. To determine whether the increased susceptibility of pre-diabetic mice to TB is reversible and is associated with dysglycemia or increased body fat mass, we performed a diet reversal experiment. Pre-diabetic mice were fed a NCD for 10 additional weeks (HFD/NCD) at which point glucose tolerance was restored, but body fat mass remained higher compared to control mice that consumed NCD throughout the entire experiment (NCD/NCD). Upon Mtb infection HFD/NCD mice had significantly lower bacterial burden compared to NCD/NCD mice and this was accompanied by restored IFN-γ responses. Our findings demonstrate that pre-diabetes increases susceptibility to TB, but a high body mass index without dysglycemia is protective. This murine model offers the opportunity to further study the underlying immunological, metabolic and endocrine mechanisms of this association.

## 1 Introduction

Tuberculosis (TB) remains one of the top 10 causes of death worldwide killing more than 1.4 million people in 2019 (WHO, 2020). Type 2 diabetes (T2D) increases the risk of developing TB as well as the risk of adverse TB treatment outcomes (Critchley et al., 2017). People with TB and T2D comorbidity have a 88% higher risk of death during treatment, a 64% higher risk of relapse and are twice as likely to develop drug-resistant TB (Huangfu et al., 2019). Paradoxically, obesity in absence of dysglycemia protects against TB (Lonnroth et al., 2010;Lin et al., 2018) and individuals with high BMI are less likely to die during TB treatment (Yen et al., 2016).

Increased susceptibility of T2D patients to TB has been attributed to poor glycemic control (Critchley et al., 2018). However, immune dysfunction and altered immunity to TB has also been demonstrated in individuals with pre-diabetes (Kumar et al., 2014;Eckold et al., 2020). Strikingly, blood transcriptomic profiles of TB patients with pre-diabetes are more similar to TB patients with T2D than those without any form of dysglycemia (Eckold et al., 2020). Whether pre-diabetes increases susceptibility to TB and TB disease severity remains unknown.

Several different animal models of TB and type 1 or type 2 diabetes have been established to study the underlying immunological mechanisms of diabetes-induced increased susceptibility to TB (Yamashiro et al., 2005;Martens et al., 2007;Sugawara and Mizuno, 2008;Vallerskog et al., 2010;Podell et al., 2014;Martinez et al., 2016;Tripathi et al., 2019;Alim et al., 2020). Such animal models mimic clinical observations from individuals with TB and diabetes co-morbidity and are particularly useful to study immune responses at the site of infection, the lung, which is difficult to achieve in patients. Given the high global prevalence rates of pre-diabetes in TB household contacts from both low and high TB burden countries - with 23% and 25% in South Africa and South Texas, respectively (Restrepo et al., 2018) - it is imperative to expand TB and diabetes association studies to include pre-diabetes. No animal models of pre-diabetes and TB have been described to date.

Here, we developed a murine high fat-diet (HFD)-induced model of pre-diabetes that presented with increased body fat mass combined with impaired glucose tolerance and hyperinsulinemia. Pre-diabetic mice had more severe TB and dysregulated cytokine production both at the site of infection and in the periphery. In subsequent diet reversal experiments, we uncoupled obesity from impaired glucose tolerance and demonstrated that obesity with restored glucose tolerance was associated with resistance to TB.

## 2 Materials and Methods

### 2.1 Ethics Statement

All experiments were carried out in accordance with protocols approved by the Health Sciences Animal Ethics Committee of The University of Queensland (MRI-UQ/413/17) and performed in accordance with the Australian Code of Practice for the Care and Use of Animals for Scientific Purposes.

### 2.2 Murine pre-diabetes and diet-reversal models

Six-week-old male C57BL/6 mice were housed in a conventional pathogen free environment, on a 12-hour light/dark cycle at 22°C and fed *ad libitum*. Animals were either fed a lard based-HFD for 12 weeks (HFD), which contained 43% available energy as fat (total fat: 23.50%, SF04-001, Specialty Feeds, Western Australia) or normal chow diet (NCD) with 12% available energy from fat for the same period (total fat: 4.60%, Standard rodent diet, Specialty Feeds, Western Australia). For the diet reversal experiment 12-week HFD fed animals were fed NCD diet for a further 10 weeks (HFD/NCD) while the control group continued on a NCD for the same period of time (NCD/NCD). Body weights of all mice were recorded weekly throughout the experiment. The respective diets continued until conclusion of the experiment. Mice were infected with Mtb H37Rv at week 12 or 22 as described below.

### 2.3 Oral glucose tolerance test, HbA1c and insulin measurement

At 12 or 22 weeks on the respective diets, oral glucose tolerance tests (OGTT), fasting insulin measurements and body composition analyses were performed. Mice were fasted for 5h with access to drinking water followed by oral gavage with 50 mg glucose per mouse. Blood was collected from tail veins and glucose concentrations were determined using a glucometer (Sensocard Plus, Elektronika Kft., Budapest, Hungary) before (0 min) and at 15, 30, 60, and 120 min after gavage. HbA1c was measured from total blood using the DCA Vantage analyzer (Siemens Healthcare Diagnostics Inc., Germany). Fasting insulin levels were quantified in serum using Ultra-Sensitive Mouse Insulin ELISA Kit (Crystal Chem, IL, USA) as per manufacturer’s instruction.

### 2.4. Body composition measurements and physiological monitoring

Whole body composition (fat and lean mass) was measured using a Bruker Minispec LF50H NMR instrument 7.5 MHz (Bruker Corporation, MA, USA) (Tinsley et al., 2004). A subset of five mice per group from the diet change experiment (at week 12 and week 22) were housed in single caging for 1 week for acclimatization followed by 3 days of metabolic profiling using the PhenoMaster System (TSE systems GmbH, Bad Homburg, Germany). Energy expenditures including CO_2_ production (VCO_2_) and O_2_ consumption (VO_2_) were monitored for 72h and respiratory exchange ratio (RER) was calculated VCO_2_/VO_2_. The mice were free to consume food and intake was measured. The resting energy expenditure (REE) was calculated using Weir equation (Weir, 1949).

### 2.5 Mtb infection, determination of bacterial burden and immunopathology

Mtb H_37_R_v_ was grown on Middlebrook 7H9 medium containing 0.05% Tween-80 and supplemented with 10% Middlebrook Oleic Albumin Dextrose Catalase Growth Supplement (BD Biosciences, USA) /0.2% glycerol to mid-log phase (OD_600_ 0.4 to 0.6). On the day of infection, a single cell suspension was prepared (O.D. of 0.1, equivalent to 50 million cells/ml) and placed in a nebulizer of an inhalation exposure system (Glas-Col, LLC, IN, USA) for aerosol infection of mice. Approximately 100-150 colony forming units (CFU) were deposited in the lung. Lungs, livers, spleens and blood were collected for determination of bacterial counts, pathology, RNA extractions and cytokine analysis as described below. For bacterial load determination tissues were homogenized, serially diluted in PBS and plated on 7H10 agar plates supplemented with 10% OADC/0.5% glycerol and incubated at 37°C. Bacterial colonies were counted after 2-3 weeks. Formalin-fixed lung lobe sections were stained with hematoxylin and eosin (H&E) and examined microscopically as previously described (Flores-Valdez et al., 2018).

### 2.6 RNA extraction and qRT-PCR

RNA was isolated from lung and blood using Isolate II RNA mini kit protocol (Bioline Reagents Ltd., London, UK) with slight modification. Briefly, blood cell pellet and lung lobes were homogenized in Trizol and vigorously mixed with chloroform (2.5:1) and centrifuged at 12,000 *x* g for 15 min at 4°C. The RNA in the aqueous phase was precipitated by mixing in cold 70% ethanol (1:2.5) followed by column-based RNA isolation using kit protocol including DNase treatment to remove genomic DNA contamination. Complementary DNA was synthesized using 2 μg of RNA and the Tetro cDNA synthesis kit (Bioline Reagents Ltd., London, UK) according to manufacturer’s instructions. Gene expression analysis was performed by quantitative real time PCR (qRT-PCR) with SensiFAST™ SYBR^®^ Lo-ROX Kit (Bioline Reagents Ltd., London, UK) run on the QuantStudio™ 7 Flex Real-Time PCR System (Applied Biosystems). All gene expression levels were normalized to *Hprt1* internal controls in each sample, and the fold changes were calculated using the 2^-ΔΔCT^ method. The list of primers used is given in Table S1.

### 2.7 ELISA

Lung homogenate supernatants were collected by centrifugation at 2000 *x* g at 4°C and stored at −80°C with protease inhibitor cocktail (Sigma). Quantification of TNF-α, IL-1β, IFN-β, CCL2, IFN-γ, and IL-10 were performed by ELISA according to the manufacturer’s instructions (R&D Systems).

### 2.8 Statistical Analysis

Data analyses were performed using GRAPHPAD PRISM Version 8 (GraphPad Software, Inc., La Jolla, CA). The results are expressed as the mean ± SEM. Comparisons between two groups were performed using non-parametric unpaired Mann-Whitney *U*-test. The relationship between two variables was ranked using Spearman’s rank correlation coefficient. Statistically significant differences between two groups are indicated in the figures as follows *, p < 0.05; **, p < 0.01; ***, p < 0.001; ****, p < 0.0001.

## 3 Results

### 3.1 Pre-diabetes increases TB severity

We developed a murine model of pre-diabetes and Mtb infection. C57BL/6 mice were fed HFD or NCD for a period of 12 weeks (Figure 1A). HFD-fed mice had significantly higher body weight (Figure 1B) and body fat mass but similar lean mass (Figure 1C) compared to NCD-fed mice. HFD-fed mice developed hyperinsulinemia indicative of insulin resistance (Figure 1D). Blood glucose concentrations after glucose challenge were higher at 15, 30, 60 and 120 min in HFD-fed mice (Figure 1E) with significantly higher area under the curve (AUC) in OGTTs (Figure 1F), while fasting blood glucose and glycated hemoglobin (HbA1c) were similar between NCD and HFD-fed mice (Figure S1). This phenotype of obesity combined with dysglycemia therefore mimics human pre-diabetes, which is characterized by insulin resistance and impaired glucose tolerance, but HbA1c levels below those of diabetes patients.

**Figure 1.**
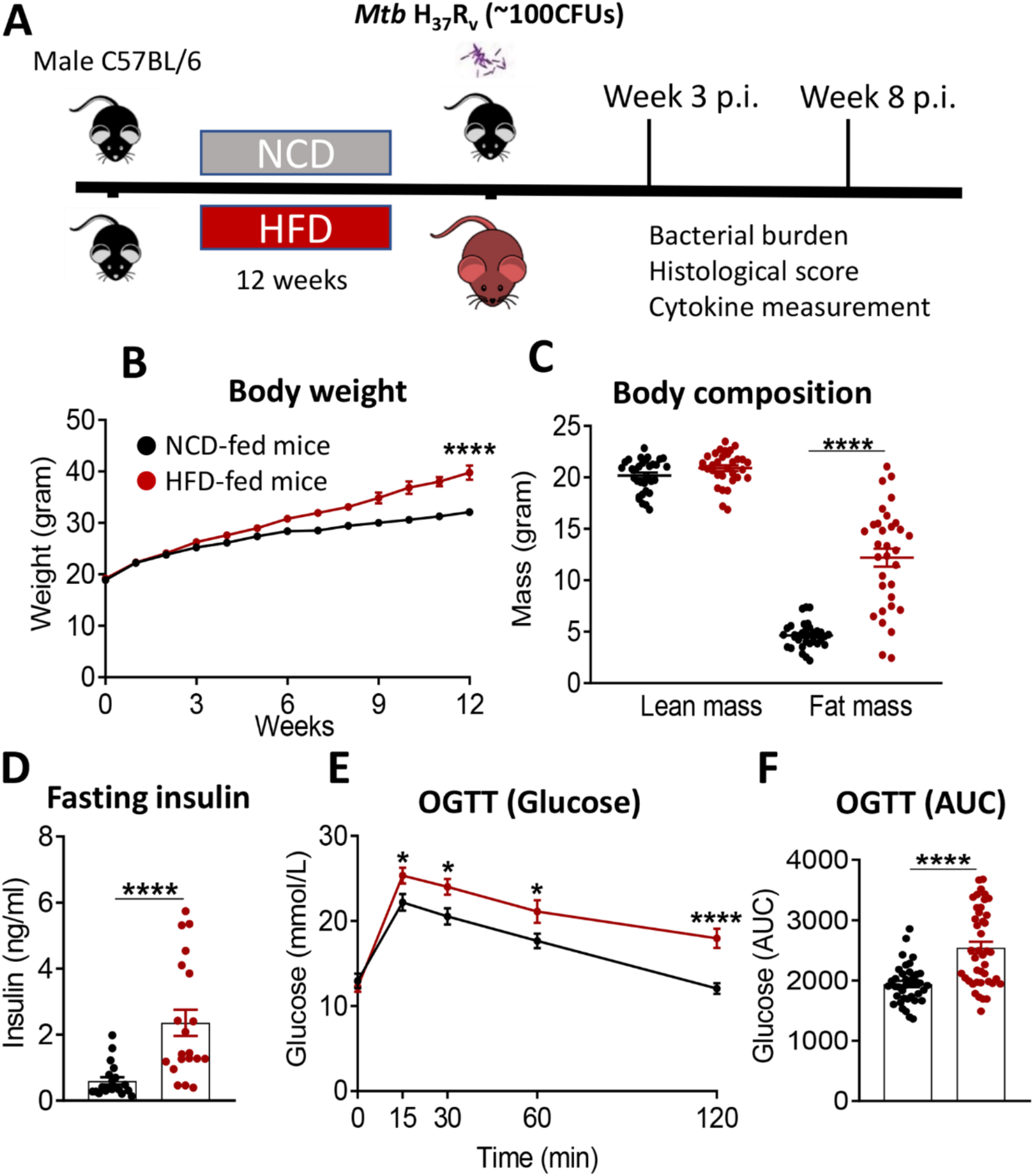
Murine model of pre-diabetes. **A.** Experimental design. **B.** Body weight of mice fed NCD (black) or HFD (red) (n=30mice/group); **C.** Body fat and lean mass at 12 weeks; **D.** fasting insulin at 12 weeks (n=20 mice/group); **E.** Blood glucose concentrations at baseline, 15, 30, 60, 120 minutes after oral glucose administration; **F.** OGTT Area under curve (AUC) (n=30 mice/group). Data are means ± SEM. Data analysis was performed by Mann-Whitney *U* test. *p < 0.05 and ****p<0.0001.

We subsequently infected the mice with live Mtb H_37_R_v_. At 3 weeks post infection (p.i.), mice with pre-diabetes had higher lung Mtb burden, although this did not reach significance (p = 0.07, Figure 2A). Lung necrosis appeared at 3 weeks p.i. in HFD-fed mice while necrosis was not detectable at this early timepoint in any NCD-fed mice. Necrosis scores were significantly higher in pre-diabetic mice by 8 weeks p.i. (Figure 2B,C) demonstrating increased lung immunopathology associated with pre-diabetes. Bacterial loads in spleens were comparable between pre-diabetic and control mice (Figure 2D). Interestingly, the Mtb burden in the fatty livers of HFD-fed mice was significantly lower than in NCD-fed mice at 3 weeks p.i, and this trend continued at 8 weeks p.i. (Figure 2E). Our data demonstrate that obesity with impaired glucose tolerance below the threshold of diabetes, i.e., pre-diabetes, increases susceptibility to pulmonary TB in a murine model.

**Figure 2.**
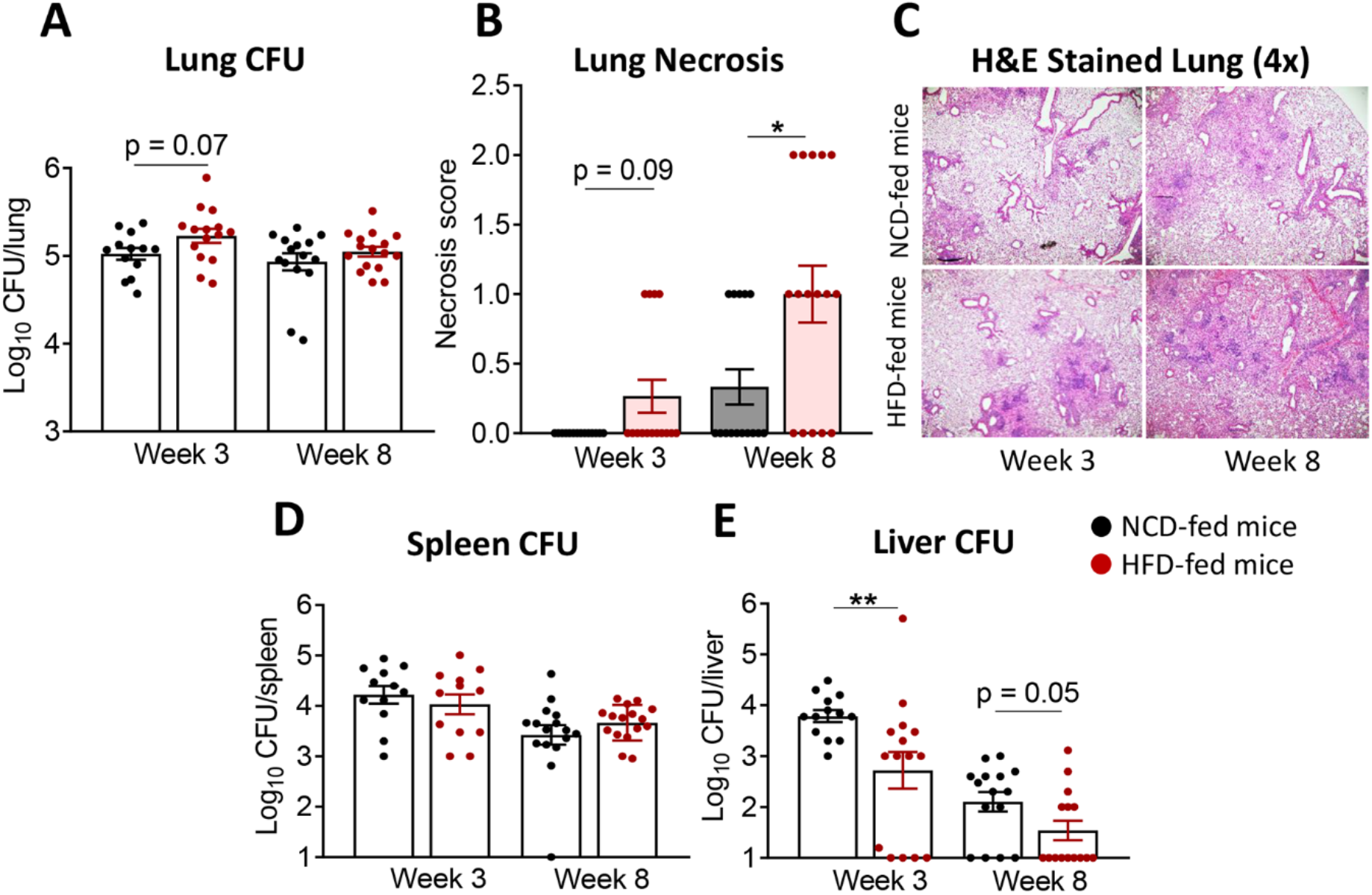
Pre-diabetes increases lung pathology in Mtb-infected mice. **A.** Lung Mtb burden in NCD-(black) and HFD-fed mice (red) at 3- and 8-weeks p.i.; **B.** Lung Necrosis scores; **C**. Representative lung histological images at 4x magnification; **D.** Mtb burden in spleen and **E.** in liver. Data are means ± SEM (n=13-16 mice/group) analyzed cumulatively in two independent experiments. Data analysis was performed by Mann-Whitney *U* test. ns = not significant *p < 0.05, and **p < 0.01.

### 3.2 Pre-diabetes alters the immune response to Mtb in the lung

Next, we investigated whether immune responses to Mtb at the site of infection are modified by pre-diabetes. We determined relative cytokine and chemokine mRNA expression and protein concentrations in lung homogenates from obese mice with impaired glucose tolerance and healthy control animals at 3 and 8 weeks p.i.. The mRNA expression of *Tnf, Ifng, Il1b, Ifnb1* and *Il10* was similar between animals (Figure 3A-E), however, mRNA expression of the chemokine *Ccl2* was significantly lower in pre-diabetic mice compared to controls at 8 weeks p.i. (Figure 3F). At the protein level, TNF-a, IFN-g and IL-10 were significantly lower at 3 weeks p.i. in pre-diabetic mice compared to control animals (Figure 3A,B,D) and concentrations of TNF-a and IFN-g remained lower also at 8 weeks p.i.. IFN-β was significantly lower in HFD-fed mice compared to NCD-fed mice at 8 weeks p.i. (Figure 3E), but IL-1β and CCL2 were not different between the groups (Figure 3C,F). In uninfected animals, no significant differences in cytokine transcript levels between NCD- and HFD-fed mice were observed at both 3-and-8 weeks p.i. (Figure S2). A low IFN-g/IL-10 ratio is a biomarker for increased TB disease severity in humans (Jamil et al., 2007) and this ratio was significantly lower in pre-diabetic mice compared to control animals at 8 weeks p.i. (Figure 3G). As expected, higher lung concentrations of TNF-α, IFN-γ, IL-1β, and CCL2 were associated with lower Mtb burden in NCD-fed mice, while this relationship was surprisingly inversed in pre-diabetic animals (Figure S3).

**Figure 3.**
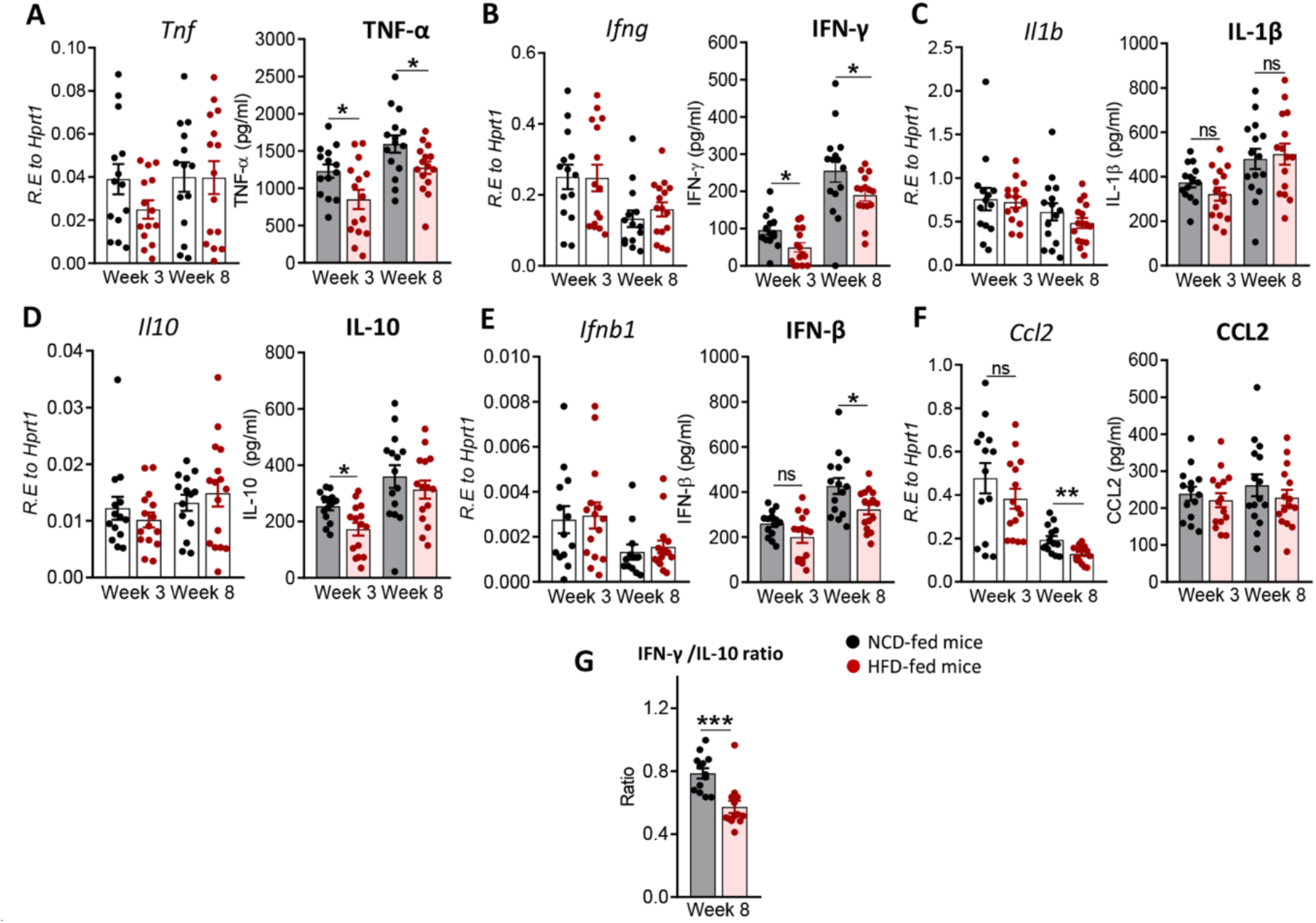
Pre-diabetes alters cytokine production in Mtb-infected lungs. Cytokine mRNA and protein levels were determined in lung homogenates by qRT-PCR and ELISA. Lung mRNA expression and protein concentrations of **A.** *Tnf*, TNF-α; **B.** *Ifng*, IFN-γ; **C.** *Il1b*, IL-1β; **D.** *Il10*, IL-10; **E.** *Ifnb1*, IFN-β and **F.** *Ccl2*, CCL2 from NCD- and HFD fed-mice at 3-and-8 weeks p.i.. **G.** IFN-γ/IL-10 ratio was determined for each mouse at week 8. Data are means ± SEM of n=13-15 mice/group analyzed cumulatively across two independent experiments. Data analysis was performed by Mann-Whitney *U* test. ns = not significant, *p < 0.05, and ***p < 0.001.

### 3.3 Pre-diabetes alters the immune response to Mtb in the periphery

To determine whether changes in the immune response to Mtb infection in pre-diabetes are limited to the site of infection or also occur in the periphery, we assessed cytokine and chemokine expression in blood from HFD- and NCD-fed mice. We found that at 3 weeks p.i. *Tnf* was higher in HFD-fed mice, but this did not reach statistical significance (p = 0.06, Figure 4A). *Il1b* was significantly lower (Figure 4C), while *Il10* was significantly higher in pre-diabetic mice compared to control animals (Figure 4D). At 8 weeks p.i. *Ifng, Il1b* and *Ccl2* were significantly lower in HFD-fed mice (Figure 4B,C,F) and we did not observe any differences in *Ifnb1* expression in HFD- *vs*. NCD-fed animals (Figure 4E). At baseline, uninfected HFD-fed mice had reduced blood transcript levels of *Ifng, Il1b, Ccl2* but increased *Il10* compared to NCD-fed mice at both 3-and-8 weeks p.i. (Figure S4). These data demonstrate that pre-diabetes-induced changes in the immune response to Mtb are not confined to the lung and occur also in the periphery.

**Figure 4.**
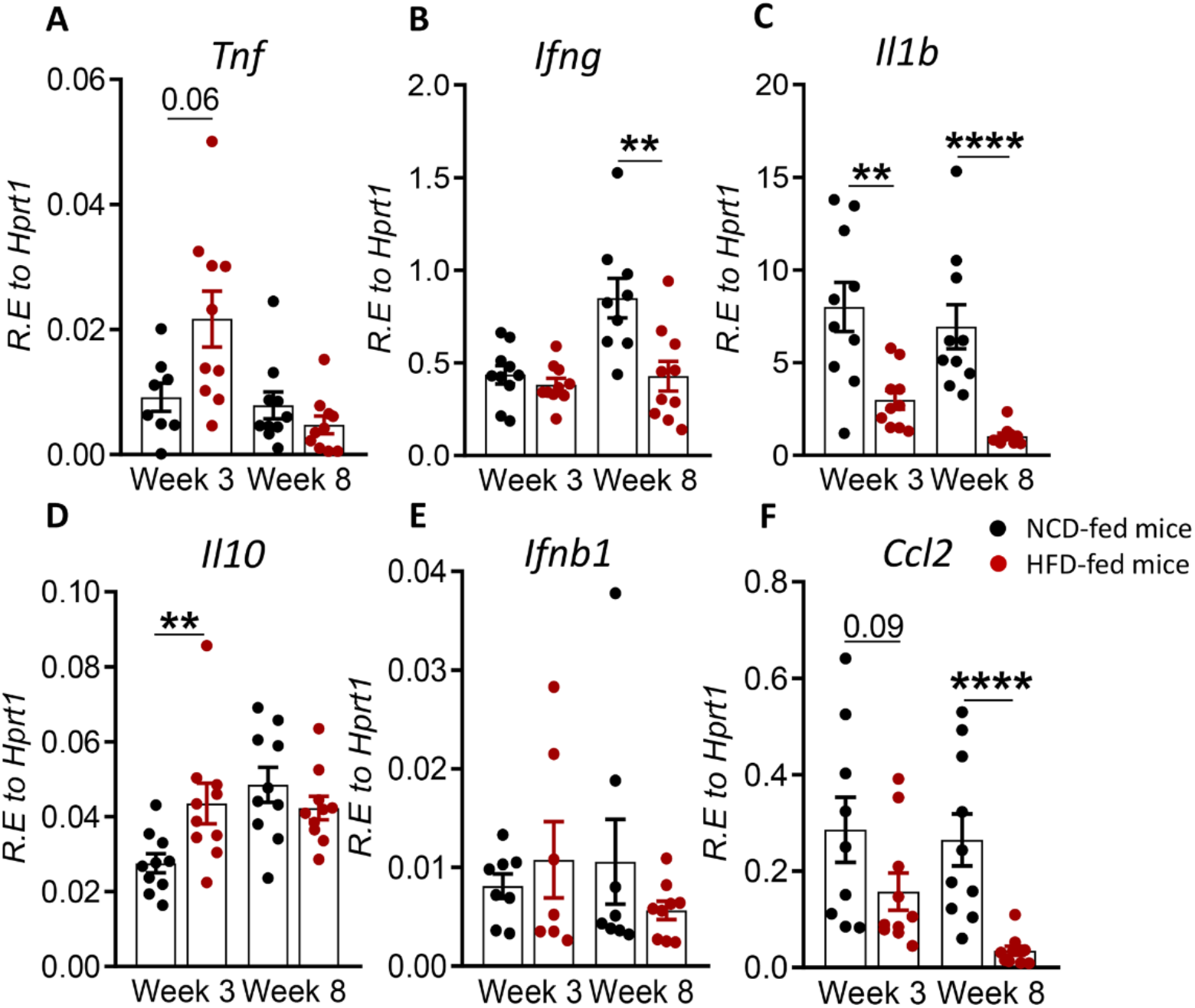
Pre-diabetes alters cytokine responses to Mtb in the periphery. Cytokine mRNA expression was determined by qRT*-*PCR in blood from NCD- and HFD-fed mice at 3- and 8-weeks p.i. **A.** *Tnf*, **B.** *Ifng*, **C.** *Il1b*, **D.** *Il10*, **E.** *Ifnb1*, and **F.** *Ccl2*. Data are means ± SEM (n=7-10 mice/group analyzed in one independent experiment). Data analysis was performed by Mann-Whitney *U* test. ns = not significant **p < 0.01, and ****p < 0.0001.

### 3.4 Restored glucose tolerance with elevated body fat mass confers mild resistance to TB

In order to assess whether a change in diet can restore glucose tolerance and reverse susceptibility to TB, we established a diet reversal model. Mice fed HFD for 12 weeks (as described for the experiment above) were subsequently fed NCD for an additional 10 weeks, here referred to as HFD/NCD mice. Control mice (NCD/NCD) were fed NCD for the entire 22 weeks (Figure 5A). Diet reversal resulted in significant loss of total body weight (Figure 5B) and fat mass (Figure 5C) in HFD/NCD animals, while the lean mass increased over time (Figure 5D). However, HFD/NCD mice maintained significantly higher body weight (Figure 5B) and higher body fat mass (Figure 5C) compared to NCD/NCD mice. Diet reversal restored the average RER and decreased the REE across 24h of light/dark phases observed in NCD-fed mice (Figure S5). Most importantly, diet reversal resulted in complete restoration of glucose tolerance between HFD/NCD and NCD/NCD animals (Figure 5E,G). We next assessed the impact of this metabolic phenotype of restored glucose tolerance but elevated body fat mass on susceptibility to TB.

**Figure 5.**
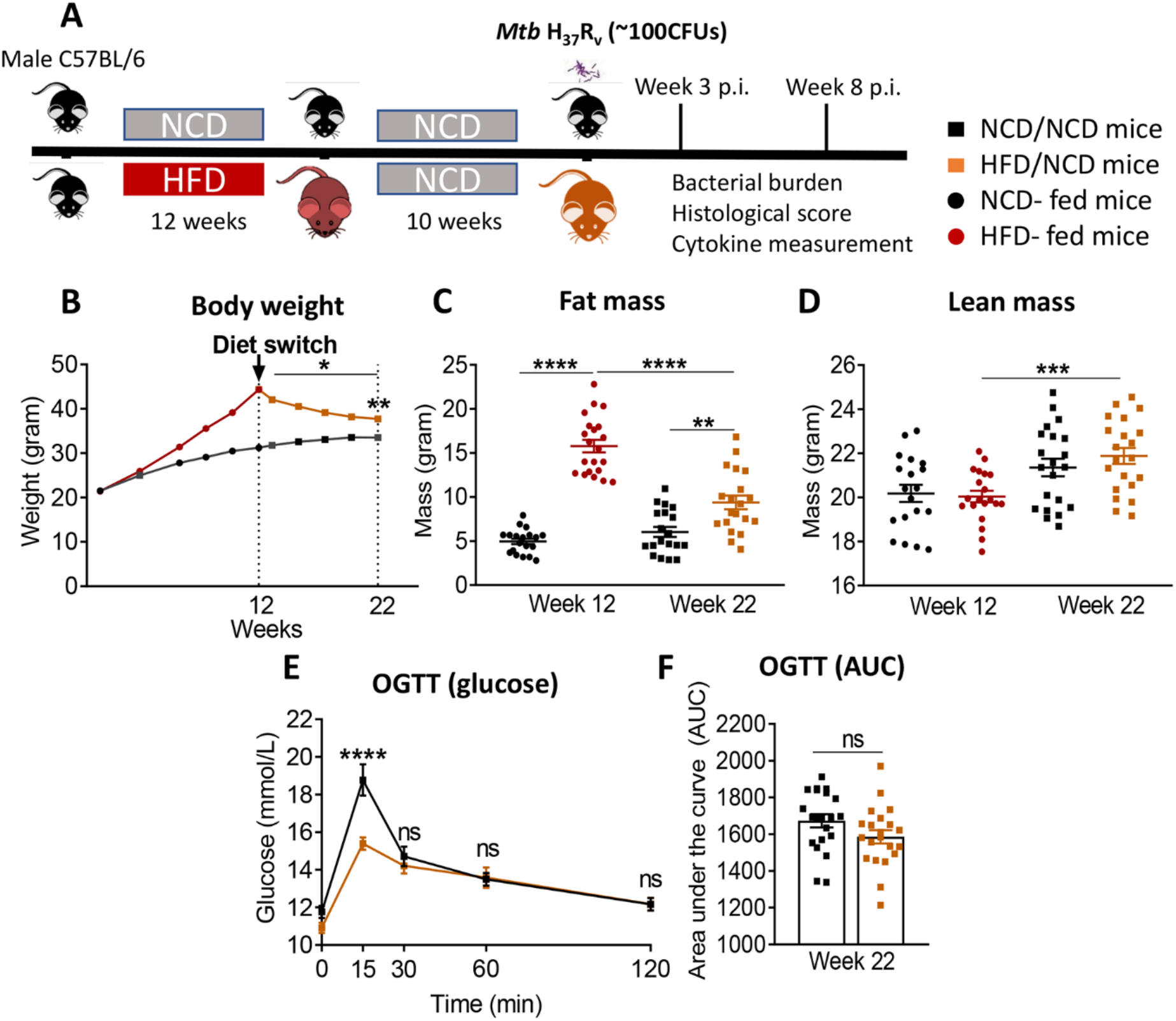
Diet reversal restores glucose tolerance while maintaining higher fat mass. **A.** Schematic showing experimental plan for diet reversal-TB model development. **B.** Body weight of mice were monitored up to 22 weeks of diet; **C.** Fat mass and **D.** lean mass were measured. OGTT was performed on NCD/NCD and HFD/NCD mice; **E.** Blood glucose concentrations at baseline, 15, 30, 60, 120 minutes after oral glucose administration and **F.** Area under curve (AUC). Data represents mean ± SEM, (n=19-20 mice/group). Data analysis was performed by Mann-Whitney *U* test. ns=not significant, *p < 0.05, **p < 0.01, ***p < 0.001 and ****p<0.0001.

HFD/NCD and NCD/NCD mice were infected with Mtb H_37_R_v_ and Mtb burden was determined at 3 and 8 weeks p.i.. Interestingly, the metabolic phenotype induced by the diet reversal, i.e., restored glucose tolerance with increased body fat mass compared to controls, conferred a mild but statistically significant resistance to TB. HFD/NCD mice had significantly lower lung Mtb burden at 8 weeks p.i. compared to control animals that only consumed NCD throughout the entire experiment (Figure 6A), while Mtb burden was similar at 3 weeks p.i.. No significant differences were observed in lung necrosis scores or spleen and liver CFU (Figure 6B-E). Coinciding with significantly reduced Mtb burden in the lung at 8 weeks p.i., we observed more foamy macrophages in lung sections from normoglycemic, obese mice compared to control animals (Figure 7) which was not evident in dysglycemic obese mice (Figure S7).

**Figure 6.**
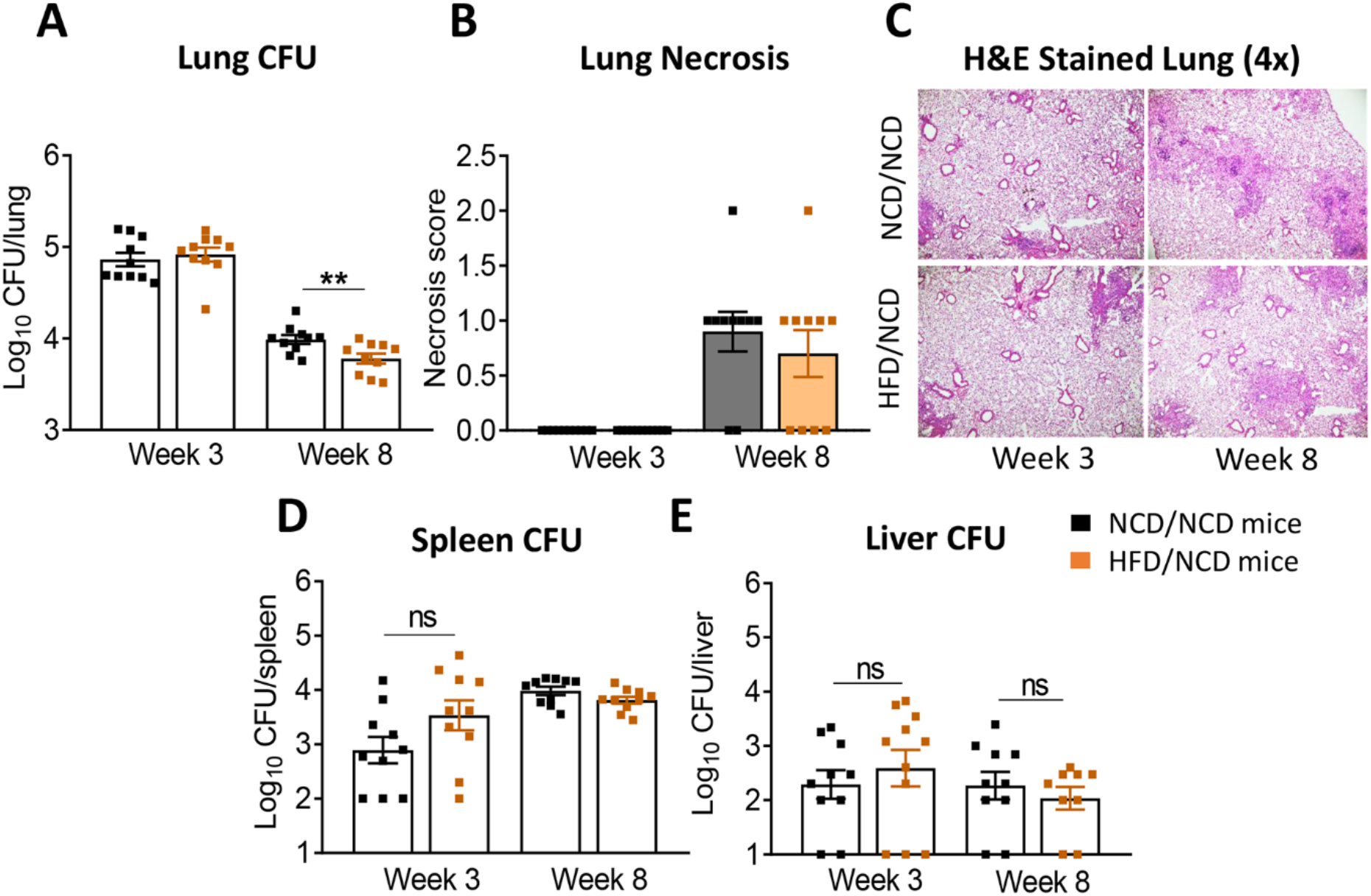
Restoration of glucose tolerance with elevated fat mass confers mild resistance to TB. **A.** Lung Mtb burden in NCD/NCD- (black) and HFD/NCD mice (orange) at 3- and 8-weeks p.i.; **B.** Lung Necrosis scores; **C.** Representative lung histological images; **D.** Mtb burden in spleen and **E.** in liver. Data are means ± SEM (n=10 mice/group analyzed in one independent experiment). Data analysis was performed by Mann-Whitney *U* test. ns = not significant.

**Figure 7.**
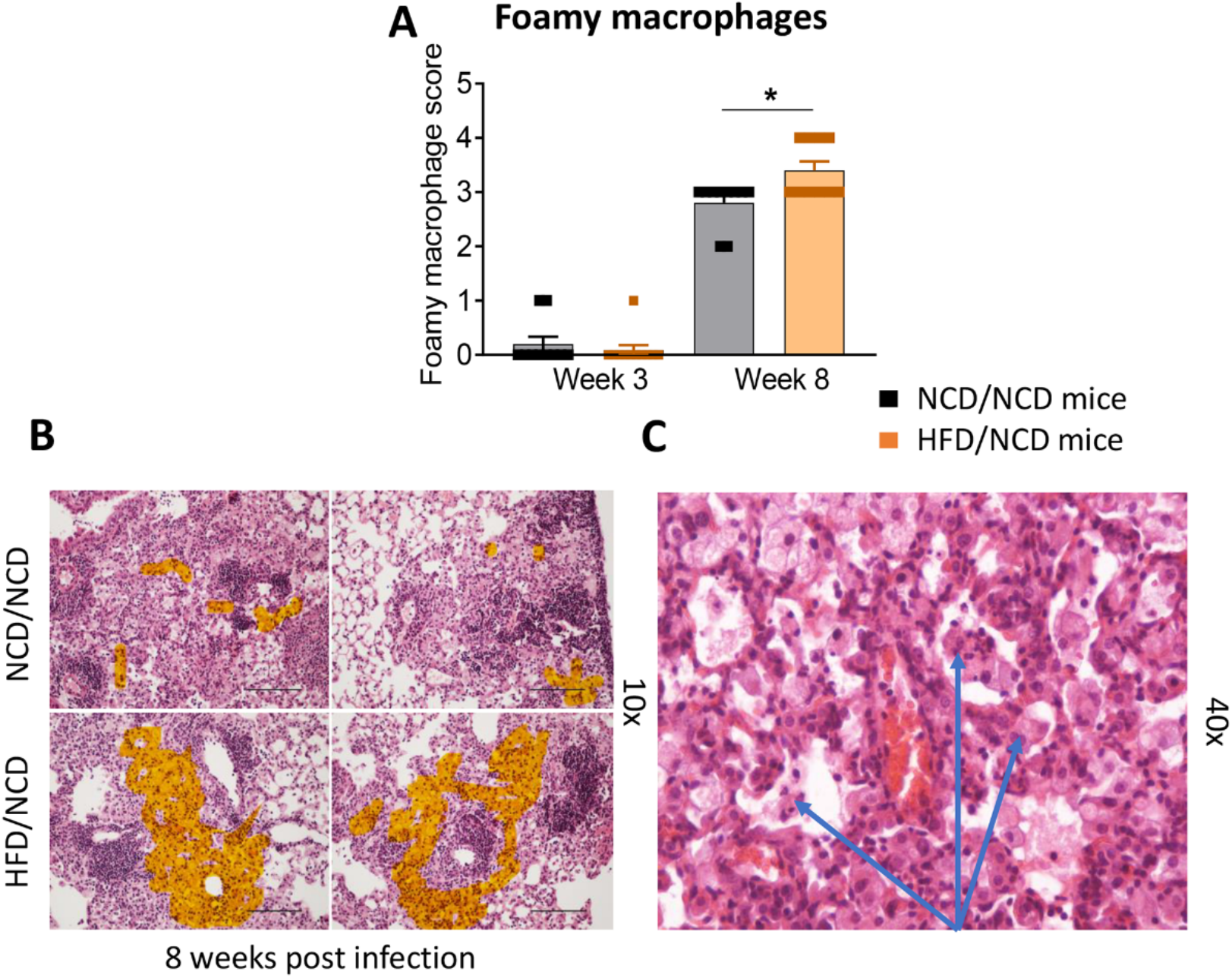
Foamy macrophages were upregulated in the lung of HFD/NCD mice. **A.** Histological scoring of foamy macrophages in lung sections of NCD/NCD mice and HFD/NCD mice at 3-and-8 weeks p.i.. **B.** Representative histological images with highlighted area (orange) of foamy macrophages; **C.** Snapshot of foamy macrophages on H&E lung sections (arrows). Data represents mean ± SEM (n=10 mice/group analyzed in one independent experiment). Data analysis was performed by Mann-Whitney *U* test. *p < 0.05.

### 3.5 Restoration of glucose tolerance improves immune responses to Mtb in the lung

We next assessed whether the change in diet and restoration of glucose tolerance while maintaining high body fat mass improves host protective immune responses to Mtb in the lung. While lung TNF-a and IFN-g concentrations were significantly lower in pre-diabetic mice compared to control mice at both 3 and 8 weeks p.i. (Figure 3A, B), concentrations of these cytokines were similar in mice with restored glucose tolerance (HFD/NCD) and their respective controls (NCD/NCD) at 8 weeks p.i., although TNF-a concentrations were still lower at 3 weeks p.i. (Figure 8A,B). At the mRNA level *Tnf, Ifng* and *Il1b* were lower in HFD/NCD *vs* NCD/NCD animals (Figure 8A,B,C). This demonstrates that production of these key cytokines for protective immune responses against Mtb was restored at the protein level by the change in diet. Similarly, IL-10 production was significantly lower in HFD-fed mice compared to NCD-fed mice at 3 weeks p.i. (Figure 3D), but after diet reversal IL-10 concentrations were comparable between HFD/NCD and NCD/NCD animals (Figure 8D). IL-1β and CCL2 concentrations, which were similar in pre-diabetic and control mice (Figure 3C,F), were significantly lower in HFD/NCD mice compared to NCD/NCD animals at 3 weeks p.i. (Figure 8C,F). While IFN-β concentrations were lower in pre-diabetic mice at 8 weeks p.i. (Figure 3E), they were lower in HFD/NCD fed mice compared to control animals at 3 weeks p.i. (Figure 8E). Correlation analysis of cytokine concentrations and lung Mtb burden are shown in Figure S6. Most importantly, the IFN-g/IL-10 ratio, which was significantly lower in pre-diabetic *vs*. control mice (Figure 3G), was now similar in animals with restored glucose tolerance and their controls (Figure 8G).

**Figure 8.**
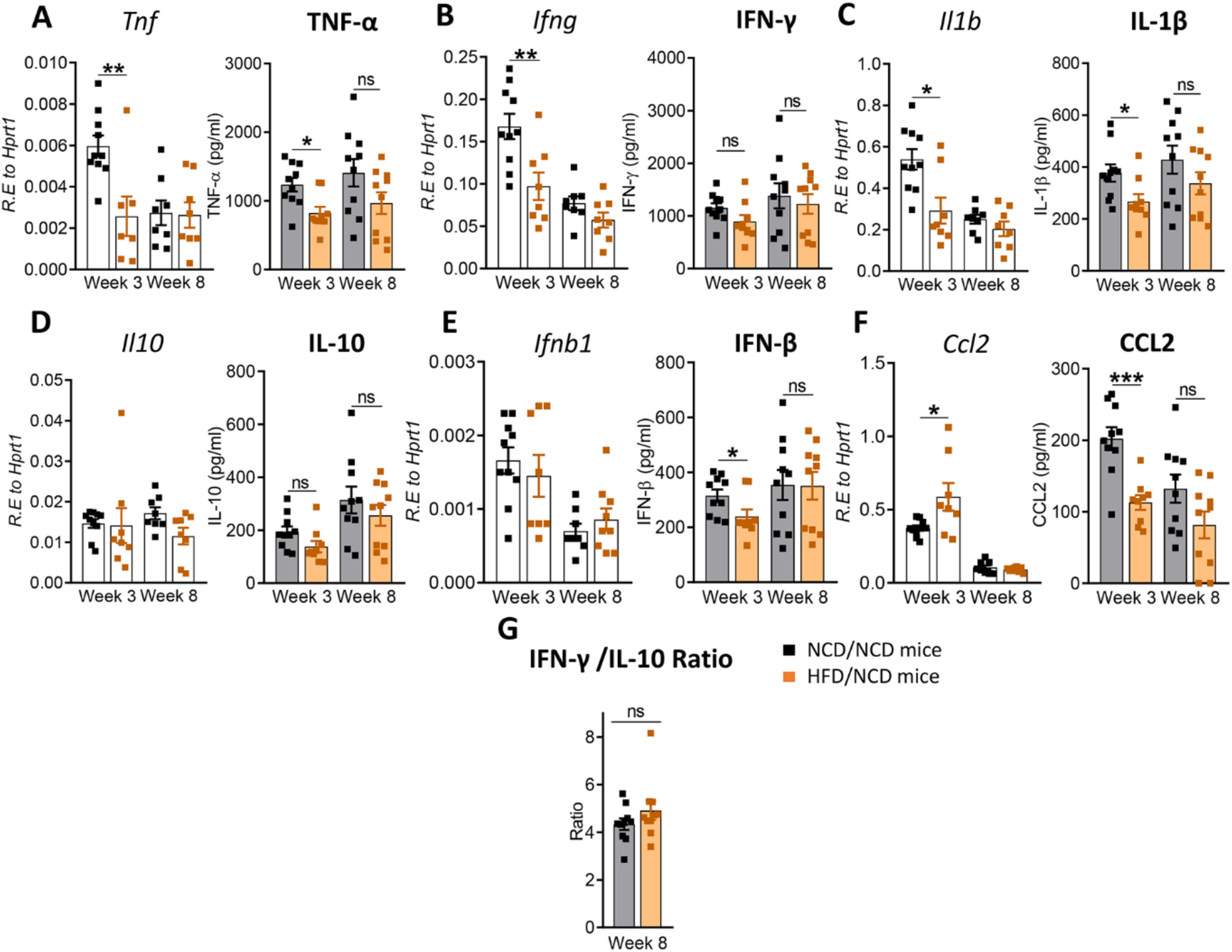
Restoration of glucose tolerance improves lung cytokine profiles. Cytokine mRNA and protein levels were determined in lung homogenates by qPCR and ELISA. Lung mRNA expression and protein concentrations of **A.** *Tnf*, TNF-α; **B.** *Ifng*, IFN-γ; **C.** *Il1b*, IL-1β; **D.** *Il10*, IL-10; **E.** *Ifnb1*, IFN-β; and **F.** *Ccl2*, CCL2 from NCD- and HFD fed-mice at 3- and 8-weeks p.i.. **G.** IFN-γ/IL-10 ratio was determined for each mouse at week 8. Data are means ± SEM (n=8-10 mice/group analyzed in one independent experiment). Data analysis was performed by Mann-Whitney *U* test. ns = not significant *p < 0.05, ** p<0.01 and ***p < 0.001.

These data demonstrate that diet reversal significantly improves this biomarker of TB disease severity.

### 3.6 Restoration of glucose tolerance improves immune responses to Mtb in the periphery

After diet reversal we also found that cytokine mRNA expression in whole blood was restored to those in control animals. For instance, mRNA expression of *Ifng, Il1b* and *Ccl2* which were lower in prediabetic mice *vs*. controls at week 8 (Figure 4B,C,F), but were similar in HFD/NCD vs. NCD/NCD animals (Figure 9B,C,F). *Il10* expression was higher in pre-diabetic animals at 3 weeks p.i. (Figure 4D) and was not significantly different in HFD/NCD *vs*. NCD/NCD animals (Figure 9D). Interestingly, blood *Ifnb* mRNA expression was significantly higher in obese mice with restored glucose tolerance compared to controls at 8 weeks p.i. (Figure 9E). Together these data demonstrate that HFD significantly impacts immune responses to Mtb at the site of infection, the lung, as well as the periphery and thus can contribute to TB disease severity.

**Figure 9.**
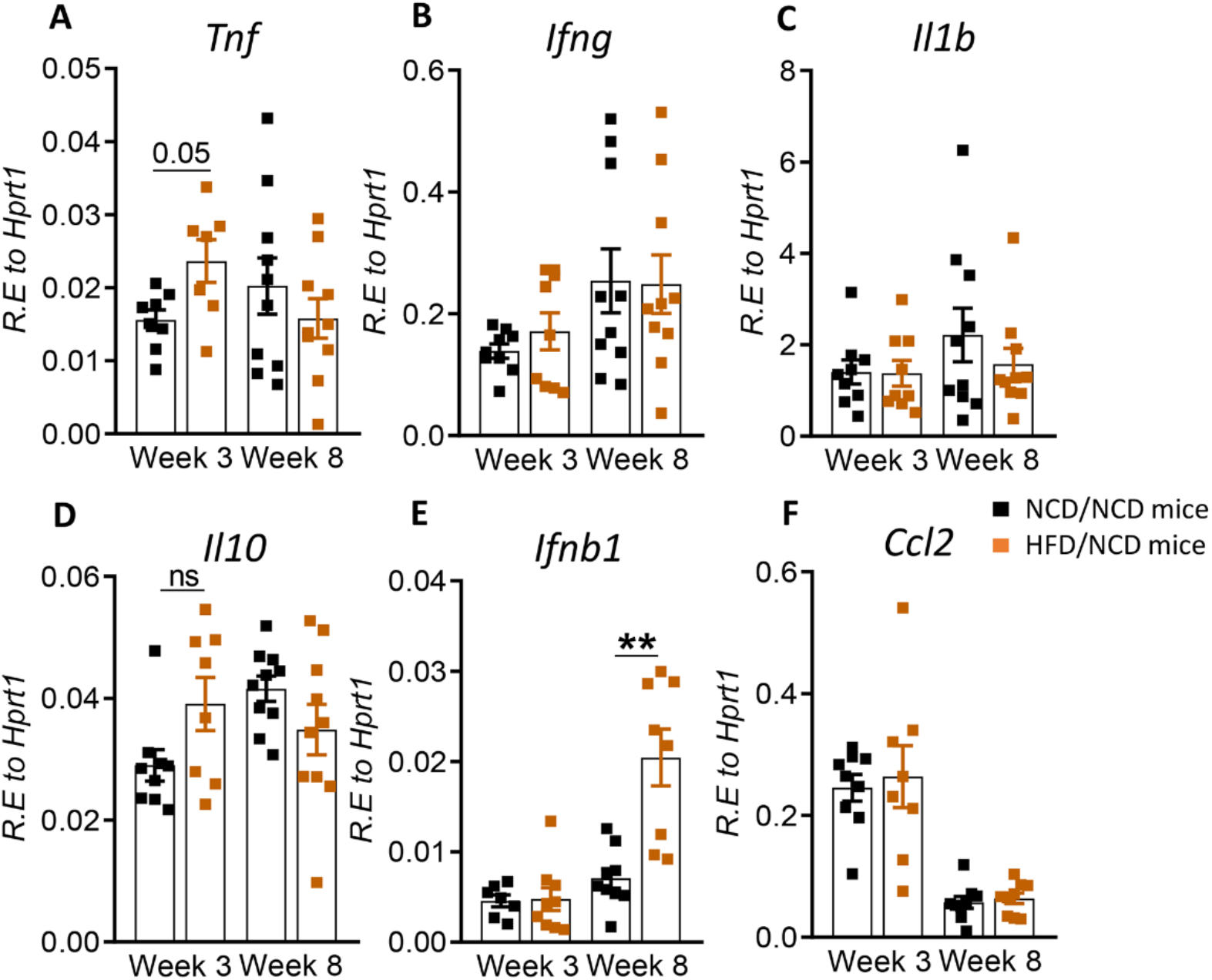
Restoration of glucose tolerance improves blood cytokine profiles. Cytokine mRNA expression was determined by qRT-PCR in blood from NCD- and HFD-fed mice at 3- and 8-weeks p.i. **A.** *Tnf*, **B.** *Ifng*, **C.** *Il1b*, **D.** *Il10*, **E.** *Ifnb1*, and **F.** *Ccl2*. Data are means ± SEM (n=8-10 mice/group analyzed in one independent experiment). Data analysis was performed by Mann-Whitney *U* test. ns = not significant **p < 0.01.

## 4. Discussion

Increased susceptibility of T2D patients to TB is well established, however, whether pre-diabetes also predisposes to more severe manifestations of pulmonary TB remains elusive as large population-based studies on the association of pre-diabetes and TB have not been performed to date. Blood transcriptomic signatures from TB patients with pre-diabetes are more similar to those from TB patients with T2D compared to TB patients without any form of dysglycemia (Eckold et al., 2020). This suggests that impaired immune responses to Mtb occur already during the early stages of dysglycemia in pre-diabetes. Given the high prevalence rates of pre-diabetes in TB endemic countries, with 27% of TB contacts in India (Shivakumar et al., 2018) and 25% in South Africa (Restrepo et al., 2018) having impaired glucose tolerance, it is imperative to investigate any associations between pre-diabetes and susceptibility to TB. To address this current knowledge gap, we developed a pre-diabetes model of Mtb infection and demonstrated more severe TB disease and altered immune responses to Mtb in the lung and blood of mice with impaired glucose tolerance.

Several different animal models of diabetes and TB exist and generally show, similarly to our prediabetes murine model, more severe disease and impaired immune responses in hyperglycemic hosts upon Mtb infection (Yamashiro et al., 2005;Martens et al., 2007;Sugawara and Mizuno, 2008;Vallerskog et al., 2010;Podell et al., 2014;Martinez et al., 2016;Tripathi et al., 2019;Alim et al., 2020). Many of these models use Streptozotocin (STZ) to induce hyperglycemia, which does not accurately reflect the chronic inflammation and vascular complications associated with human T2D. Nevertheless, these models provide valuable insight into hyperglycemia associated immune impairment. STZ-induced chronic hyperglycemia resulted in increased Mtb lung burden, more inflammation and lower IFN-g production in the lung (Martens et al., 2007). Similarly, we found lower IFN-g and TNF-a production in the lungs of pre-diabetic mice combined with more severe immunopathology and significantly lower IFN-g/IL-10 ratios, a biomarker for TB disease severity (Jamil et al., 2007). The increased susceptibility of STZ-treated mice was attributed to a delayed innate immune response due to impaired recognition of Mtb by alveolar macrophages from hyperglycemic animals, which subsequently results in delayed adaptive immune responses (Martinez et al., 2016). It is likely that pre-diabetic mice also have a delayed adaptive immune response given the lower IFN-production, however, whether this is due to impaired recognition of Mtb by pre-diabetic alveolar macrophages remains to be elucidated in future studies. Vallerskog *et al*. reported lower CCL2 expression in the lungs of STZ-induced hyperglycemic mice (Vallerskog et al., 2010). While prediabetic animals in our model showed lower *Ccl2* mRNA at 8-weeks p.i., protein concentrations of CCL2 were not significantly different. Eckhold *et al*. found reduced type I IFN responses in blood transcriptomic signatures from TB patients with pre-diabetes (Eckold et al., 2020). We did not observe reduced *Ifnb1* mRNA expression in blood, however, IFN-β concentrations were significantly reduced in lungs of pre-diabetic mice at 8 weeks p.i.. A HFD-based model of T2D and TB recently demonstrated moderately higher Mtb burden in the early stages of infection at 2 weeks, but not during late infection, and reduced IFN-g production in HFD-fed diabetic mice (Alim et al., 2020), which is consistent with our data in this pre-diabetes model. An interesting observation in our murine prediabetes model was the significantly reduced Mtb burden in the liver of HFD-fed animals. This finding is in line with human studies showing that diabetes does not increase the risk of developing extrapulmonary TB (Magee et al., 2016) despite a higher risk of pulmonary TB. Increased hepatic Mtb burden was however observed in the HFD-based murine model of diabetes (Alim et al., 2020), but in this study the animals were infected *via* the intra-venous route and not via the natural aerosol route, which likely explain the increased hepatic Mtb burden. HFD-induced alterations in gut microflora leads to severe pulmonary damage and mortality in Toll-like receptor deficient mice (Ji et al., 2014) and it is possible that dysbiosis of the gut microbiota contributes in part to susceptibility of our pre-diabetic animals to TB.

To determine whether the increased susceptibility of pre-diabetic mice is due to impaired glucose tolerance or obesity, we performed a diet reversal experiment in which we could separate impaired glucose tolerance from obesity. Surprisingly, obese animals with restored glucose tolerance were able to contain Mtb better than their healthy-weight controls. In these animals more lung macrophages had a foamy macrophage phenotype. It is possible that these macrophages contained Mtb in a quiescent non-replicating state resulting in overall lower Mtb burden (Russell et al., 2009). The restoration of glucose tolerance, while maintaining a high body fat mass, also resulted in restoration of IFN-g responses. CCL2 concentrations on the other hand were significantly lower compared to NCD-fed mice. This may serve as a feedback mechanism to limit further recruitment of macrophages to the lung. The change in diet ultimately improved the IFN-γ/IL-10 ratio and necrosis scores were similar in obese animals with a history of glucose impairment compared to healthy chow-fed animals. Observations from our murine model are consistent with findings in humans where obesity in absence of dysglycemia protects against TB (Lonnroth et al., 2010;Lin et al., 2018).

Therefore, both our HFD-induced pre-diabetes model and the diet reversal model of Mtb infection mimic observations in humans. Our murine models offer the unique opportunity to elucidate the underlying immune-metabolic mechanisms of obesity-induced resistance *vs*. dysglycemia-associated susceptibility to TB. Importantly, our data provide clear evidence, that immune impairment to Mtb including decreased lung IFN-g production indicative of delayed adaptive immune priming occurs already during pre-diabetes and likely contributes to more severe disease. Thus, large population-based studies are warranted to determine the impact of pre-diabetes in susceptibility to TB in TB-endemic countries.

## Author Contributions

RS, MDN and KR wrote the manuscript; RS, MDN, SK, MLD, JK, AB carried out the experiments, RS, MDN, SB, HBO analyzed the data and compiled the figures; RS, MDN, SB, HBO, SK, SH, AB, CC, KS and KR interpreted the data and contributed intellectually.

## Acknowledgments

We thank Adrian T. Gemiarto for technical assistance and the staff of the biological resource facility at the Translational Research Institute for animal husbandry.

## Data Availability Statement

The raw data supporting conclusions of this article will be made available by the authors, without undue reservation.

## Supplementary Material

**Figure S1.**
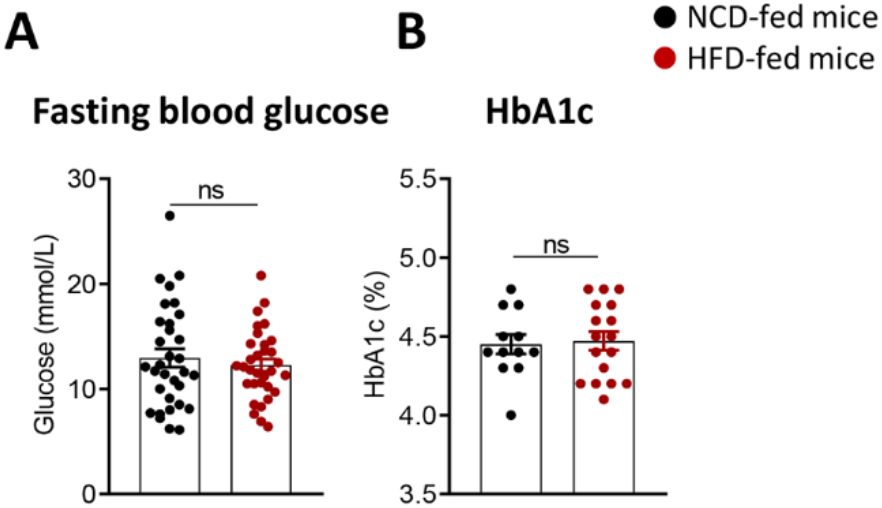
Characterization of pre-diabetes in C57BL/6 mice. **A.** Fasting blood glucose was measured after 12 weeks of NCD or HFD (n=30 mice/group analyzed cumulatively across two independent experiments). **B.** Glycated hemoglobin (HbA1c) was measured in whole blood on a randomly selected subset of mice from each group (n=12-17 mice/group analyzed cumulatively across two independent experiments). Data analysis was performed by Mann-Whitney *U* test. ns = not significant.

**Figure S2.**
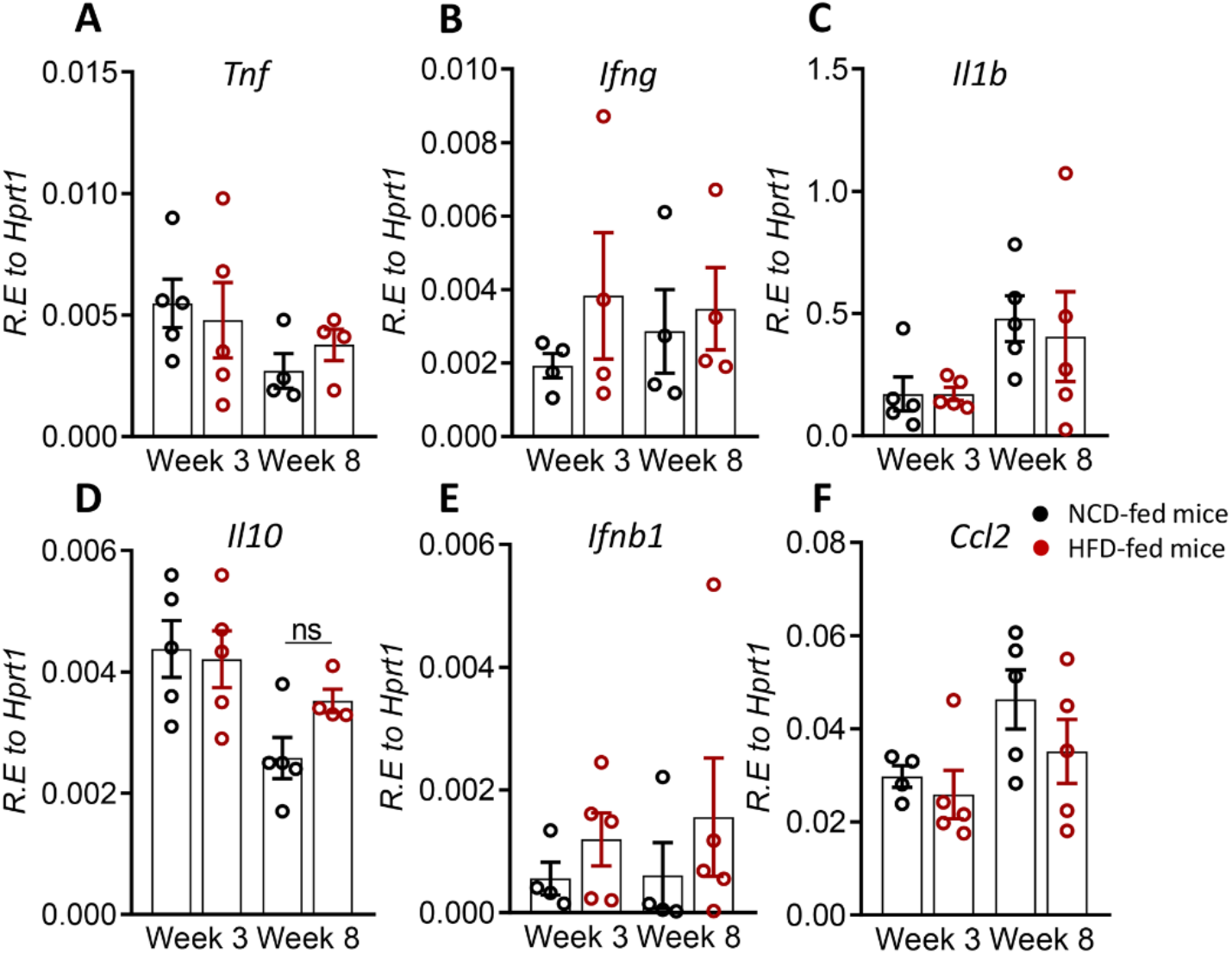
Inflammatory cytokine mRNA expression in the lung from uninfected NCD- and HFD-fed mice. Relative expression of **A.** *Tnf*, **B.** *Ifng*, **C.** *Il1b*, **D.** *Il10*, **E.** *Ifnb1* and **F.** *Ccl2*. Data are means ± SEM of n=4-5 mice/group analyzed in one independent experiment. Data analysis was performed by Mann-Whitney *U* test. ns = not significant.

**Figure S3.**
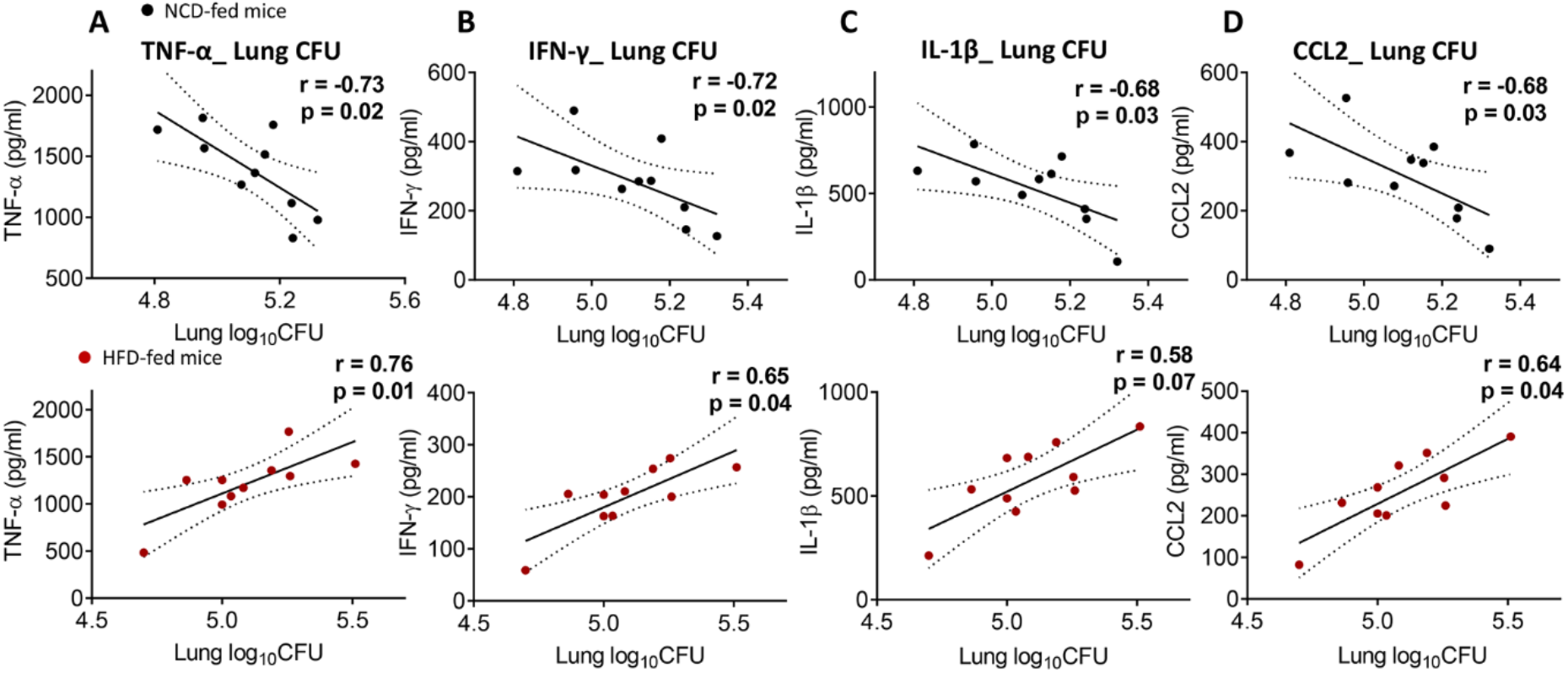
Correlation between lung bacterial burden and lung pro-inflammatory cytokines/chemokines in NCD-fed mice and HFD-fed mice. **A.** TNF-α, **B.** IFN-γ, **C.** IL-1β and **D.** CCL2 concentrations in NCD or HFD mice correlated with their respective CFUs using the Spearman’s rank correlation test. (n=10 mice/group). Spearman r and respective p values have been shown on the figure.

**Figure S4.**
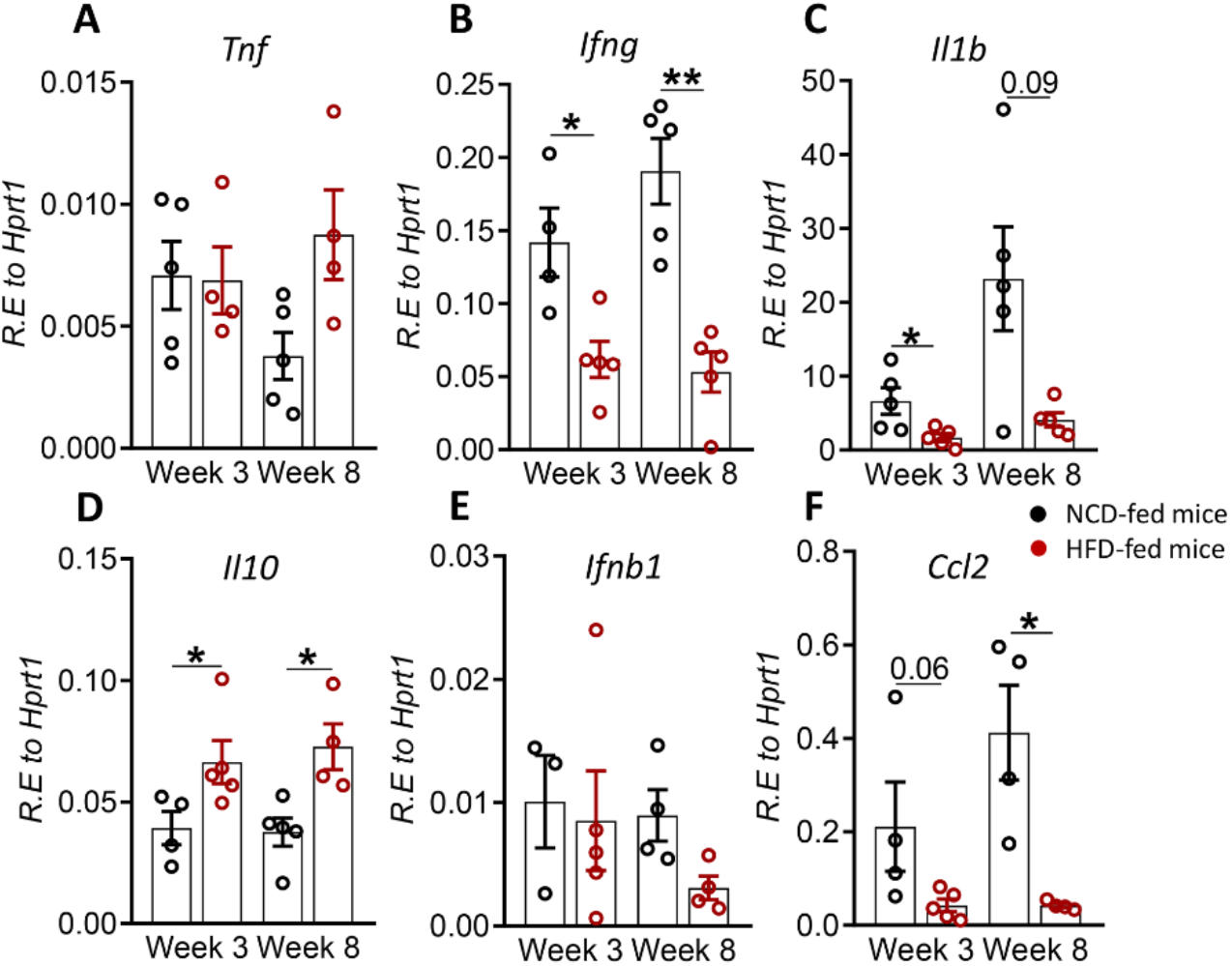
mRNA expression of inflammatory cytokines in blood from uninfected NCD- and HFD-fed mice. Relative expression of **A.** *Tnf*, **B.** *Ifng*, **C.** *Il1b*, **D.** *Il10*, **E.** *Ifnb1* and **F.** *Ccl2*. Data are means ± SEM of n=4-5 mice/group analyzed in one independent experiment. Data analysis was performed by Mann-Whitney *U* test. *p < 0.05, **p < 0.01.

**Figure S5.**
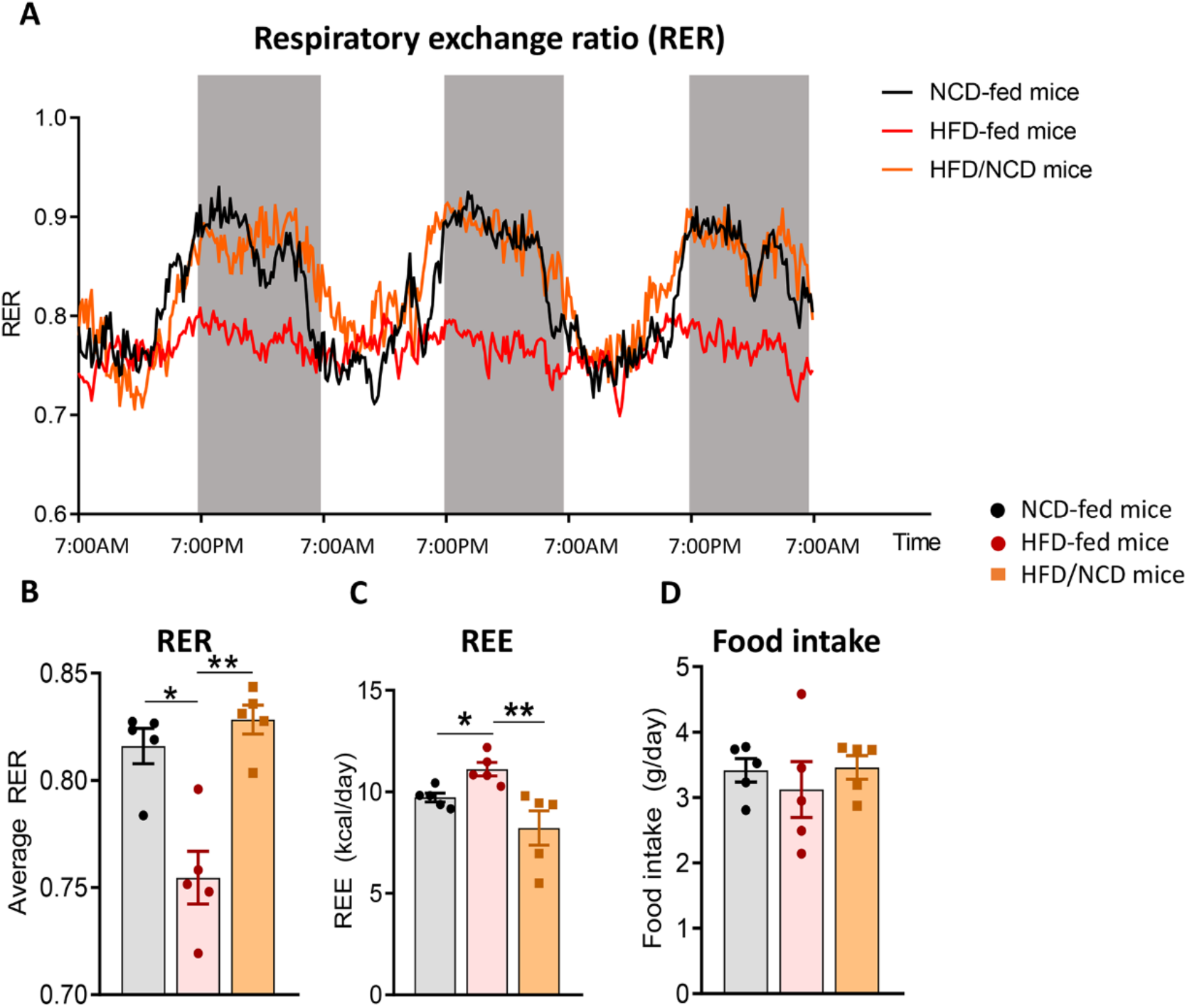
Metabolic profile of NCD-, HFD- and HFD/NCD mice. **A.** Changes in RER throughout light/dark phases. Average daily values of **B.** RER, **C.** REE and **D.** food intake. Data are shown as mean ± SEM. (n=5 mice/group). *P < 0.05, **P < 0.01.

**Figure S6.**
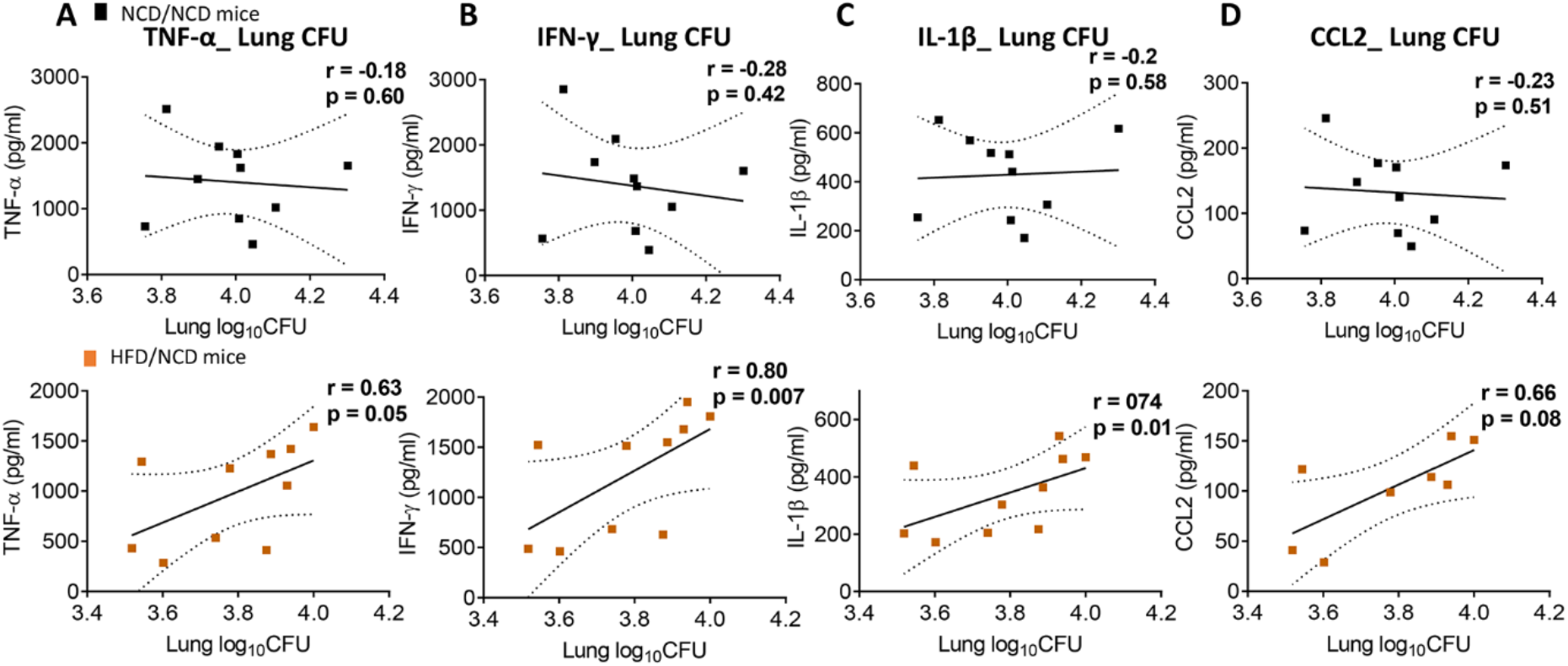
Correlation between lung bacterial burden and lung pro-inflammatory cytokines/chemokines in NCD/NCD and HFD/NCD mice. **A.** TNF-α, **B.** IFN-γ, **C.** IL-1β and **D.** CCL2 concentrations correlated with CFUs using the Spearman’s rank correlation test. (n=10 mice/group). Spearman r and respective p values have been shown on the figure.

**Figure S7.**
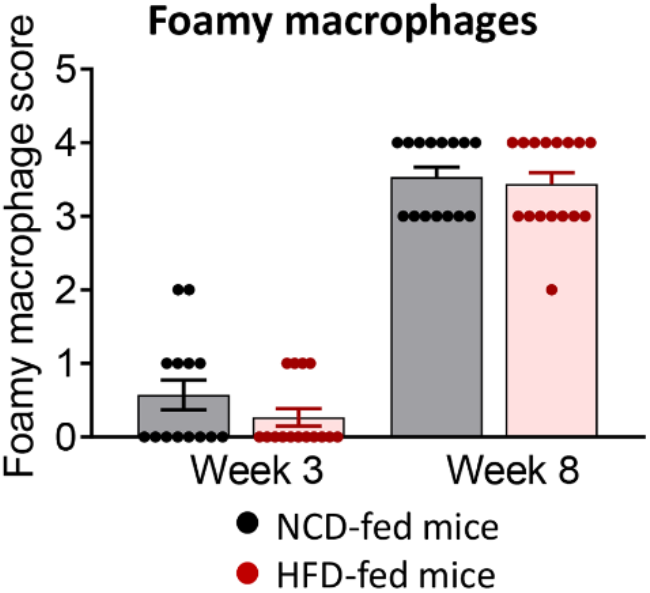
Foamy macrophages in lungs from NCD- and HFD-fed mice. Histological scoring of appearance of foamy macrophages in lung sections from infected mice at 3 weeks and 8 weeks postinfection. Data represents mean ± SEM (n=10 mice/group analyzed in one independent experiment). Data analysis was performed by Mann-Whitney *U* test.

**Table S1.**
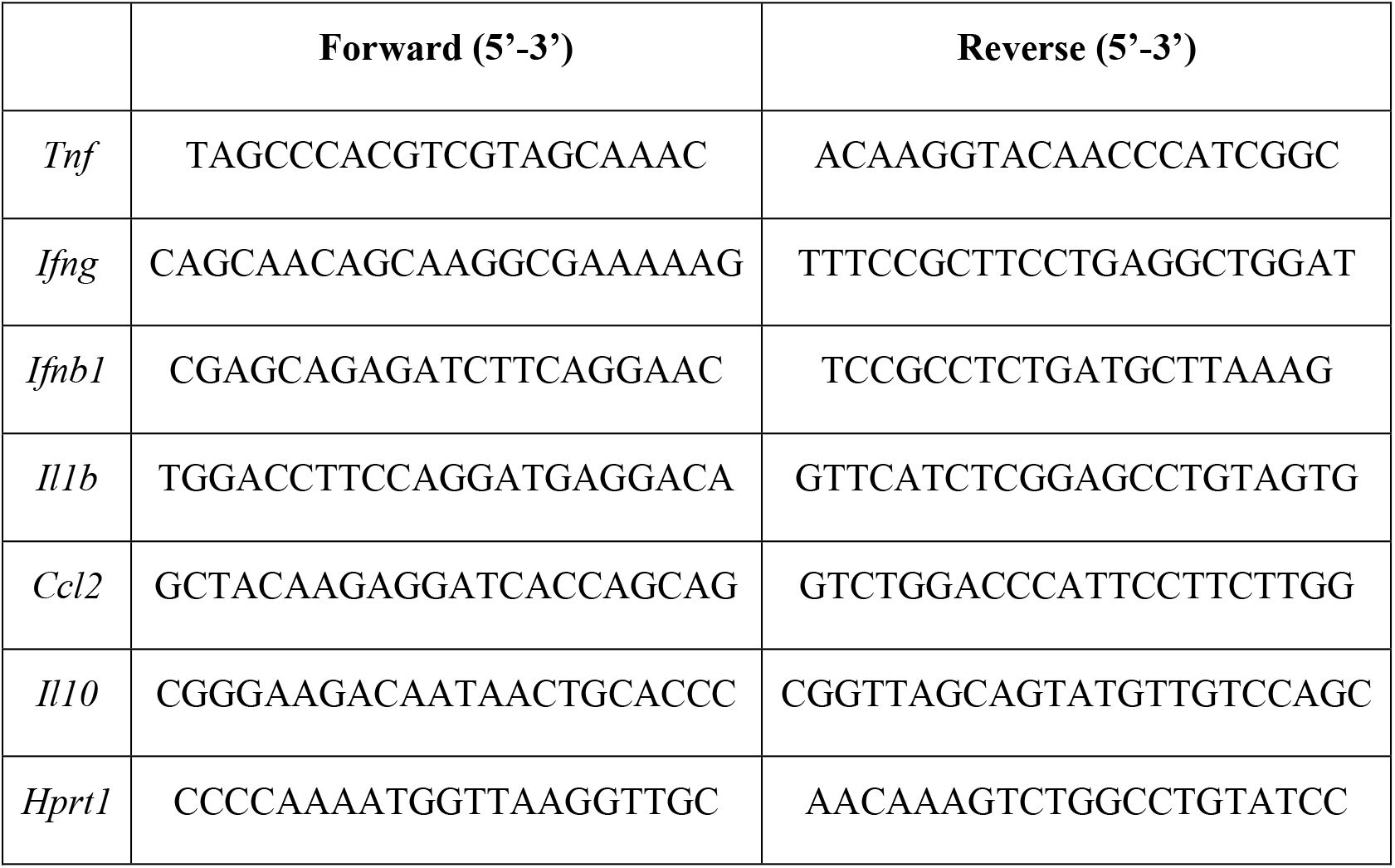
List of mouse primer sequences.

## References

Alim, M.A., Kupz, A., Sikder, S., Rush, C., Govan, B., and Ketheesan, N. (2020). Increased susceptibility to Mycobacterium tuberculosis infection in a diet-induced murine model of type 2 diabetes. Microbes Infect 22, 303–311.

Critchley, J.A., Carey, I.M., Harris, T., Dewilde, S., Hosking, F.J., and Cook, D.G. (2018). Glycemic Control and Risk of Infections Among People With Type 1 or Type 2 Diabetes in a Large Primary Care Cohort Study. Diabetes Care 41, 2127–2135.

Critchley, J.A., Restrepo, B.I., Ronacher, K., Kapur, A., Bremer, A.A., Schlesinger, L.S., Basaraba, R., Kornfeld, H., and Van Crevel, R. (2017). Defining a Research Agenda to Address the Converging Epidemics of Tuberculosis and Diabetes: Part 1: Epidemiology and Clinical Management. Chest 152, 165–173.

Eckold, C., Kumar, V., Weiner Rd, J., Alisjahbana, B., Riza, A.L., Ronacher, K., Coronel, J., Kerry-Barnard, S., Malherbe, S.T., Kleynhans, L., Stanley, K., Ruslami, R., Ioana, M., Ugarte-Gil, C., Walzl, G., Van Crevel, R., Wijmenga, C., Critchley, J.A., Dockrell, H.M., and Cliff, J.M. (2020). Impact of intermediate hyperglycaemia as well as diabetes on immune dysfunction in tuberculosis. Clin Infect Dis.

Flores-Valdez, M.A., Pedroza-Roldan, C., Aceves-Sanchez, M.J., Peterson, E.J.R., Baliga, N.S., Hernandez-Pando, R., Troudt, J., Creissen, E., Izzo, L., Bielefeldt-Ohmann, H., Bickett, T., and Izzo, A.A. (2018). The BCGDeltaBCG1419c Vaccine Candidate Reduces Lung Pathology, IL-6, TNF-alpha, and IL-10 During Chronic TB Infection. Front Microbiol 9, 1281.

Huangfu, P., Ugarte-Gil, C., Golub, J., Pearson, F., and Critchley, J. (2019). The effects of diabetes on tuberculosis treatment outcomes: an updated systematic review and meta-analysis. Int J Tuberc Lung Dis 23, 783–796.

Jamil, B., Shahid, F., Hasan, Z., Nasir, N., Razzaki, T., Dawood, G., and Hussain, R. (2007). Interferon gamma/IL10 ratio defines the disease severity in pulmonary and extra pulmonary tuberculosis. Tuberculosis (Edinb) 87, 279–287.

Ji, Y., Sun, S., Goodrich, J.K., Kim, H., Poole, A.C., Duhamel, G.E., Ley, R.E., and Qi, L. (2014). Diet-induced alterations in gut microflora contribute to lethal pulmonary damage in TLR2/TLR4-deficient mice. Cell Rep 8, 137–149.

Kumar, N.P., Banurekha, V.V., Nair, D., Sridhar, R., Kornfeld, H., Nutman, T.B., and Babu, S. (2014). Coincident pre-diabetes is associated with dysregulated cytokine responses in pulmonary tuberculosis. PLoS One 9, e112108.

Lin, H.H., Wu, C.Y., Wang, C.H., Fu, H., Lonnroth, K., Chang, Y.C., and Huang, Y.T. (2018). Association of Obesity, Diabetes, and Risk of Tuberculosis: Two Population-Based Cohorts. Clin Infect Dis 66, 699–705.

Lonnroth, K., Williams, B.G., Cegielski, P., and Dye, C. (2010). A consistent log-linear relationship between tuberculosis incidence and body mass index. Int J Epidemiol 39, 149–155.

Magee, M.J., Foote, M., Ray, S.M., Gandhi, N.R., and Kempker, R.R. (2016). Diabetes mellitus and extrapulmonary tuberculosis: site distribution and risk of mortality. Epidemiol Infect 144, 2209–2216.

Martens, G.W., Arikan, M.C., Lee, J., Ren, F., Greiner, D., and Kornfeld, H. (2007). Tuberculosis susceptibility of diabetic mice. Am J Respir Cell Mol Biol 37, 518–524.

Martinez, N., Ketheesan, N., West, K., Vallerskog, T., and Kornfeld, H. (2016). Impaired Recognition of Mycobacterium tuberculosis by Alveolar Macrophages From Diabetic Mice. J Infect Dis 214, 1629–1637.

Podell, B.K., Ackart, D.F., Obregon-Henao, A., Eck, S.P., Henao-Tamayo, M., Richardson, M., Orme, I.M., Ordway, D.J., and Basaraba, R.J. (2014). Increased severity of tuberculosis in Guinea pigs with type 2 diabetes: a model of diabetes-tuberculosis comorbidity. Am J Pathol 184, 1104–1118.

Restrepo, B.I., Kleynhans, L., Salinas, A.B., Abdelbary, B., Tshivhula, H., Aguillon-Duran, G.P., Kunsevi-Kilola, C., Salinas, G., Stanley, K., Malherbe, S.T., Maasdorp, E., Garcia-Viveros, M., Louw, I., Garcia-Oropesa, E.M., Lopez-Alvarenga, J.C., Prins, J.B., Walzl, G., Schlesinger, L.S., and Ronacher, K. (2018). Diabetes screen during tuberculosis contact investigations highlights opportunity for new diabetes diagnosis and reveals metabolic differences between ethnic groups. Tuberculosis (Edinb) 113, 10–18.

Russell, D.G., Cardona, P.J., Kim, M.J., Allain, S., and Altare, F. (2009). Foamy macrophages and the progression of the human tuberculosis granuloma. Nat Immunol 10, 943–948.

Shivakumar, S., Chandrasekaran, P., Kumar, A.M.V., Paradkar, M., Dhanasekaran, K., Suryavarshini, N., Thomas, B., Kohli, R., Thiruvengadam, K., Kulkarni, V., Hannah, L.E., Sivaramakrishnan, G.N., Pradhan, N., Dolla, C., Gupte, A., Ramachandran, G., Deluca, A., Meshram, S., Bhardawaj, R., Bollinger, R.C., Golub, J., Selvaraj, K., Gupte, N., Swaminathan, S., Mave, V., and Gupta, A. (2018). Diabetes and pre-diabetes among household contacts of tuberculosis patients in India: is it time to screen them all? Int J Tuberc Lung Dis 22, 686–694.

Sugawara, I., and Mizuno, S. (2008). Higher susceptibility of type 1 diabetic rats to Mycobacterium tuberculosis infection. Tohoku J Exp Med 216, 363–370.

Tinsley, F.C., Taicher, G.Z., and Heiman, M.L. (2004). Evaluation of a quantitative magnetic resonance method for mouse whole body composition analysis. Obes Res 12, 150–160.

Tripathi, D., Radhakrishnan, R.K., Sivangala Thandi, R., Paidipally, P., Devalraju, K.P., Neela, V.S.K., Mcallister, M.K., Samten, B., Valluri, V.L., and Vankayalapati, R. (2019). IL-22 produced by type 3 innate lymphoid cells (ILC3s) reduces the mortality of type 2 diabetes mellitus (T2DM) mice infected with Mycobacterium tuberculosis. PLoS Pathog 15, e1008140.

Vallerskog, T., Martens, G.W., and Kornfeld, H. (2010). Diabetic mice display a delayed adaptive immune response to Mycobacterium tuberculosis. J Immunol 184, 6275–6282.

Weir, J.B. (1949). New methods for calculating metabolic rate with special reference to protein metabolism. J Physiol 109, 1–9.

Who (2020). Global Tuberculosis Report 2020.

Yamashiro, S., Kawakami, K., Uezu, K., Kinjo, T., Miyagi, K., Nakamura, K., and Saito, A. (2005). Lower expression of Th1-related cytokines and inducible nitric oxide synthase in mice with streptozotocin-induced diabetes mellitus infected with Mycobacterium tuberculosis. Clin Exp Immunol 139, 57–64.

Yen, Y.F., Chuang, P.H., Yen, M.Y., Lin, S.Y., Chuang, P., Yuan, M.J., Ho, B.L., Chou, P., and Deng, C.Y. (2016). Association of Body Mass Index With Tuberculosis Mortality: A Population-Based Follow-Up Study. Medicine (Baltimore) 95, e2300.

